# The role of gene dosage in budding yeast centrosome scaling and spontaneous diploidization

**DOI:** 10.1101/2020.06.04.133975

**Authors:** Jingjing Chen, Zhiyong Xiong, Danny E. Miller, Zulin Yu, Scott McCroskey, William D. Bradford, Ann M. Cavanaugh, Sue L. Jaspersen

## Abstract

Ploidy is the number of whole sets of chromosomes in a species. Ploidy is typically a stable cellular feature that is critical for survival. Polyploidization is a route recognized to increase gene dosage, improve fitness under stressful conditions and promote evolutionary diversity. However, the mechanism of regulation and maintenance of ploidy is not well characterized. Here, we examine the spontaneous diploidization associated with mutations in components of the *Saccharomyces cerevisiae* centrosome, known as the spindle pole body (SPB). Although SPB mutants are associated with defects in spindle formation, we show that two copies of the mutant in a haploid yeast favors diploidization in some cases, leading us to speculate that the increased gene dosage in diploids ‘rescues’ SPB duplication defects, allowing cells to successfully propagate with a stable diploid karyotype. This copy number-based rescue is linked to SPB scaling: certain SPB subcomplexes do not scale or only minimally scale with ploidy. We hypothesize that acquisition of lesions in structures with incompatible allometries such as the centrosome may drive changes such as whole genome duplication, which have shaped the evolutionary landscape of many eukaryotes.

**Author Summary:** Ploidy is the number of whole sets of chromosomes in a species. Most eukaryotes alternate between a diploid (two copy) and haploid (one copy) state during their life and sexual cycle. However, as part of normal human development, specific tissues increase their DNA content. This gain of entire sets of chromosomes is known as polyploidization, and it is observed in invertebrates, plants and fungi, as well. Polyploidy is thought to improve fitness under stressful conditions and promote evolutionary diversity, but how ploidy is determined is poorly understood. Here, we use budding yeast to investigate mechanisms underlying the ploidy of wild-type cells and specific mutants that affect the centrosome, a conserved structure involved in chromosome segregation during cell division. Our work suggests that different scaling relationships (allometry) between the genome and cellular structures underlies alterations in ploidy. Furthermore, mutations in cellular structures with incompatible allometric relationships with the genome may drive genomic changes such duplications, which are underly the evolution of many species including both yeasts and humans.

## Introduction

Multiple conserved processes act together to ensure eukaryotic cells maintain a stable chromosome composition, called the karyotype. Most organisms have a diploid karyotype with two copies of each chromosome (Ezov et al. 2006). In nature, fungi are also commonly diploids, however, a haploid karyotype can be stably maintained in most lab strains (Gerstein and Otto 2009; Herskowitz 1988). Changes in the karyotype through gains or losses of one or more chromosomes leads to aneuploidy, which is associated with miscarriage, cancer and fungal drug resistance (Gianaroli et al. 1999; Storchova and Pellman 2004; Selmecki, Forche, and Berman 2006; Gordon, Resio, and Pellman 2012). Gains of whole sets of chromosomes (polyploidy) is another type of karyotype alteration that has driven evolution of many eukaryotes, including vertebrates and yeast such as *Saccharomyces cerevisiae* (Lee, Davidson, and Duronio 2009; Selmecki et al. 2015; Mittal et al. 2017; Otto 2007; Harari et al. 2018). Increased ploidy is observed in certain highly differentiated human tissues such as liver parenchyma, heart muscle, placenta and bone marrow (Anatskaya and Vinogradov 2007; Liu et al. 2010; Orr-Weaver 2015), and it is frequently observed in plants (Soltis et al. 2015). However, polyploidy is also linked to aneuploidy as increased ploidy often leads to chromosome instability (CIN) (Amend et al. 2019; Storchova and Pellman 2004; Kops, Weaver, and Cleveland 2005; Fujiwara et al. 2005; Mayer and Aguilera 1990). For example, in budding yeast the rate of chromosome loss in triploids and tetraploids is 30- and 1000-fold higher than haploids (Andalis et al. 2004). The mechanism(s) resulting in CIN in polyploids are poorly understood but may be linked to incompatible allometries (biological scaling relationships) driven by increasing genome size (Varberg and Jaspersen 2018; Hara and Kimura 2011; Storchova et al. 2006; Andalis et al. 2004).

The cell division cycle is a highly conserved process that ensures chromosomes are replicated and segregated into daughter cells. Throughout eukaryotes, chromosomes are distributed into daughter cells by the mitotic spindle, a microtubule network formed around two spindle poles known as centrosomes in metazoans or spindle pole bodies (SPBs) in fungi. Duplication of the centrosome/SPB is coupled with the cell cycle such that cells entering mitosis have exactly two spindle poles to form a bipolar spindle (Ruthnick and Schiebel 2018; Cavanaugh and Jaspersen 2017). Errors in centrosome duplication result in the formation of monopolar or multipolar spindles. This has long been considered a driving factor in aneuploidy and polyploidy despite mechanisms to cluster multipolar spindles or surveillance mechanisms to detect spindle defects (Vitre and Cleveland 2012). In *Saccharomyces cerevisiae* a mutant defective in SPB duplication was isolated by Lee Hartwell in his famous screen for cell division cycle mutants (Hartwell et al. 1973). *cdc31-1* (allelic to *cdc31-2* used here) mutants arrest in metaphase due to monopolar spindles. Although the mutant was isolated in a yeast with a haploid karyotype, *cdc31-1* cells were found to be polyploid (diploid) (Schild, Ananthaswamy, and Mortimer 1981). This phenotype of spontaneous diploidization is not unique to *cdc31-1*, also having been described for other SPB components and regulators (Sing et al. 2018; Jaspersen, Giddings, and Winey 2002; Jaspersen et al. 2006; Chen et al. 2019; Kupke et al. 2011; Vallen et al. 1994; Winey et al. 1991; Chial et al. 1999).

In the electron microscope, the SPB appears as a trilaminar plaque-like structure embedded in the nuclear membrane (Byers and Goetsch 1974). Associated with one side is a specialized region of the nuclear envelope known as the half-bridge. Molecularly, the ~1 GDa SPB is composed of 18 components present in multiple copies. Each protein localizes to a distinct region in the SPB, including the core, the half-bridge or the membrane region that anchors the SPB to the nuclear envelope (referred to here as the SPB luminal ring) (Cavanaugh and Jaspersen 2017; Ruthnick and Schiebel 2018). The half-bridge plays an essential role in SPB duplication while the ultimate function of the core is microtubule nucleation.

Polyploidy may result from errors in chromosome segregation, but, somewhat paradoxically, increases in ploidy expand the burden of chromosomes that must be replicated and segregated by the cell cycle machinery. While polyploidy does not lead to the proteotoxic stress observed in many aneuploids (Sheltzer and Amon 2011; Oromendia and Amon 2014), genetic analysis of haploid, diploid and tetraploid yeast cells pointed to three processes that are essential for genome stability in cells of higher ploidy (tetraploids) but non-essential in cells of lower ploidy (haploids and diploids): homologous recombination, sister chromatid cohesion and mitotic spindle function (Storchova et al. 2006; Storchova 2014). In yeast, where a single microtubule binds to each chromosome via its kinetochore (Joglekar et al. 2006), the number of microtubules must scale with ploidy. Consistent with this idea, the size of the SPB core, measured by electron microscopy (EM) as the diameter across its central region, increases linearly with ploidy: from 90-110 nm in haploids, to 160 nm in diploids and 240 nm in tetraploids (Adams and Kilmartin 1999; Kilmartin 2003; Byers and Goetsch 1974). How the SPB scales in size is unknown. The simplest idea, that polyploids have extra copies of SPB genes, seems insufficient as the SPB of haploid cells can also scale in size when the cell cycle is delayed or when the number of centromeres is increased (Winey et al. 1993; Nannas et al. 2014). In addition, in SPB mutants that spontaneously diploidize, the cell must build a larger SPB and nucleate more microtubules – so it is unclear why the mutation would not result in another error in segregation that would further increase ploidy.

In metazoans, centrosome size also correlates with spindle size, and changes in its size have been linked to defects in chromosome segregation, aneuploidy and cancer (Vitre and Cleveland 2012; Cabral et al. 2019; Marshall 2011; Marteil et al. 2018). Here, we used budding yeast as a model system to examine the relationship between ploidy changes and SPB size scaling at a molecular level. We examined the polyploidy associated with SPB mutants and performed a genetic screen to isolate suppressors of *cdc31-2* diploidization. We found that spontaneous diploidization rescues the growth defect associated with some, but not all, SPB mutants. Our data shows that mutations that are rescued by increased ploidy are only found in genes encoding specific SPB components that localize to regions of the SPB structure that we show do not scale linearly with chromosome number. We propose that polyploidy acts as a ‘dosage’ suppressor and that acquisition of malfunctional centrosomes could drive eukaryotic evolution or disease progression by promoting changes such as whole genome duplication.

## Results

### Spontaneous diploidization in SPB mutants

Cdc31 is the yeast centrin ortholog, a small, highly conserved calcium binding protein present at centrosomes and other microtubule-organizing centers (MTOCs) across eukaryotes. A temperature-sensitive mutation in *cdc31-2* (E133K) causes haploid yeast cells to undergo spontaneous diploidization at the permissive growth temperature (23°C) immediately upon loss of a wild-type copy of *CDC31*, a phenotype we will refer to as increase-in-ploidy (IPL) (Chan and Botstein 1993). The IPL phenotype is observed by flow cytometry as 2N and 4N peaks compared to the 1N and 2N peaks seen in haploid cells (Figure 1A). Whole genome sequencing (WGS) of *cdc31-2* mutants shows that the IPL is an example of autopolyploidy, with two exact copies of each chromosome (diploid control in Figure 2D, Table S1). Examination of spindle structure by fluorescence microscopy showed that 59% of *cdc31-2* large budded cells contained a bipolar spindle (Figure 1A). Interestingly, we did not observe evidence of *cdc31-2* progressing to tetraploids, suggesting that these mutants are maintained as stable diploids with fully compensated microtubule nucleation ability as a result of their IPL phenotype. Together, these results show that loss of Cdc31 function at 23°C can be overcome by spontaneous transition to a diploid state.

**Figure 1.**
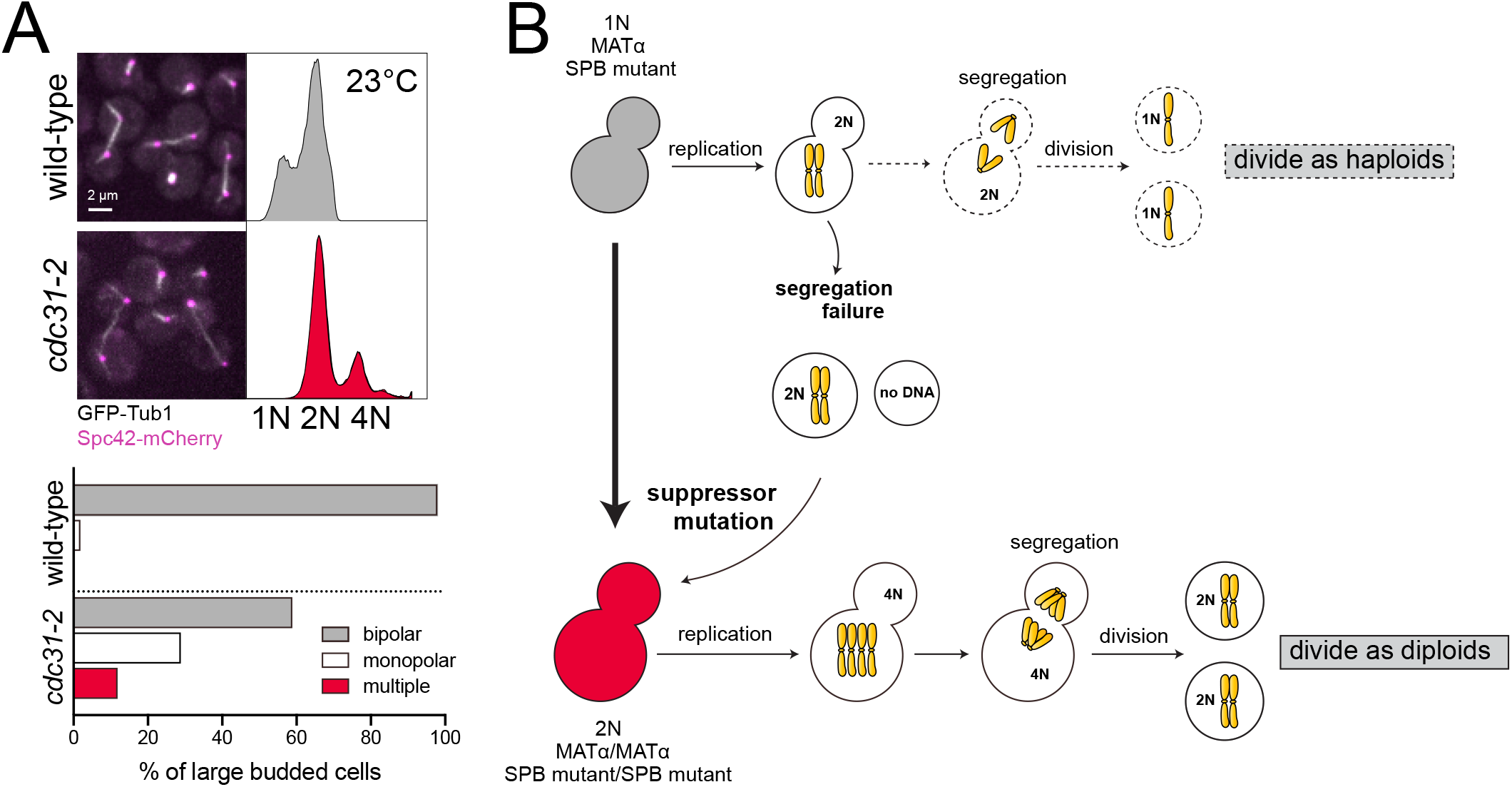
Spontaneous diploidization of SPB mutants such as *cdc31-2* at the permissive temperature. (A) Wild-type (SLJ7819) and *cdc31-2* mutant (SLJ10777) cells containing GFP-Tub1 (white) and Spc42-mCherry (magenta) were generated with a *pURA3-CDC31* plasmid. After growth on 5-FOA at 23°C to select for loss of the plasmid, the *cdc31-2* mutant spontaneously diploidize despite the formation of bipolar spindles. A representative image from each is shown (bar = 2 μm), and the percentage of large budded cells for with bipolar, monopolar or multipolar/broken spindles was quantitated (n>150). DNA content was assayed by flow cytometry. The biphasic peaks in wild-type cells represent cells with G1 (1N) and G2/M (2N) DNA content. At 23°C, *cdc31-2* mutants have diploid DNA content (2N and 4N). (B) Schematic of pathway to diploidization in *cdc31-2*. Due to a defect in chromosome segregation, haploid (1N) cells undergo an aberrant cell division to produce a diploid (2N) and aploid (0N) cell. The diploid cell does not have the same defect as haploids, resulting in successfully propagation. Because of this, we suspect that a suppressor mutation is acquired.

**Figure 2.**
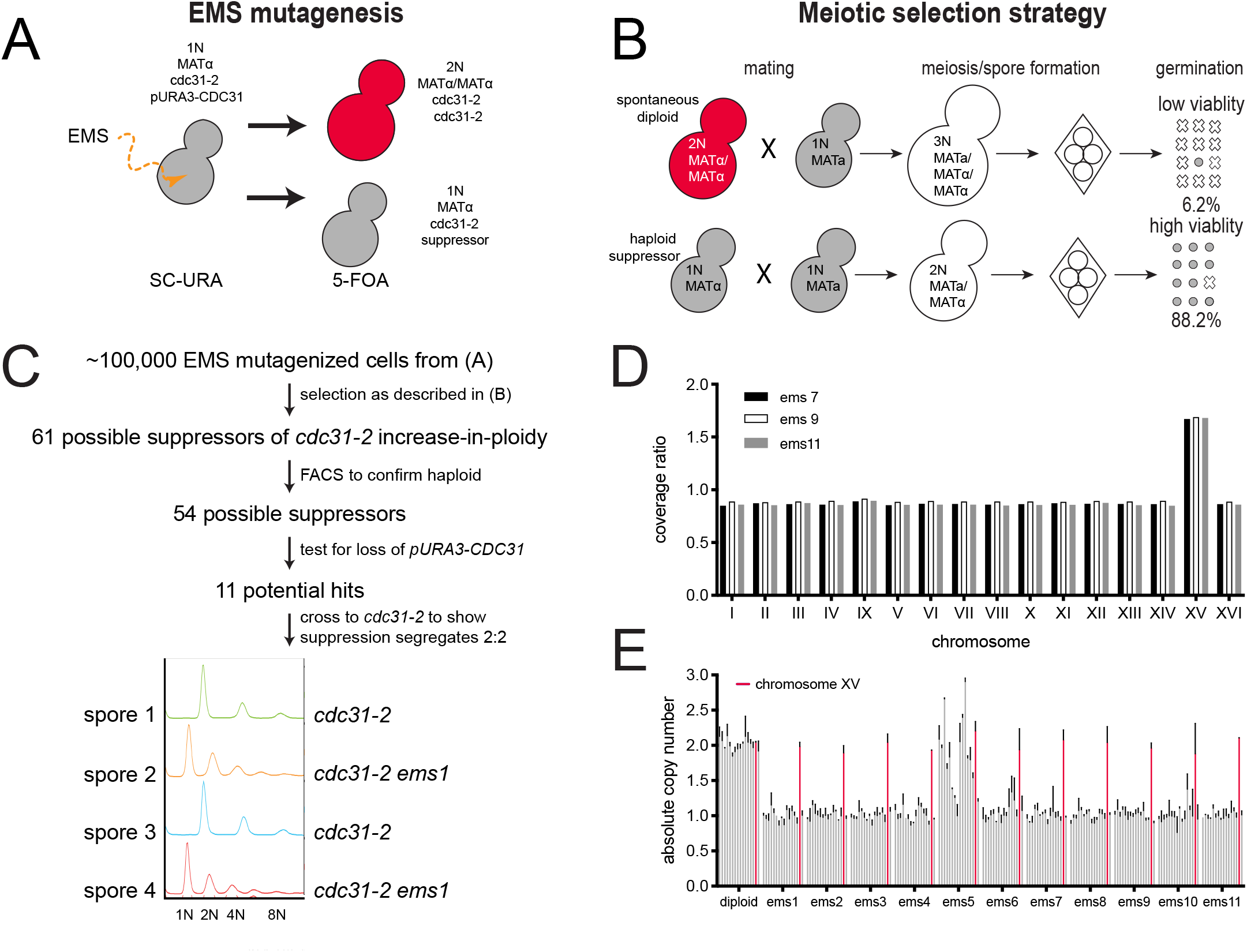
Screen for suppressors of *cdc31-2* diploidization. (A) Suppressors of the *cdc31-2* increase-in-ploidy were isolated following mutagenesis of SLJ6749 (*MATα cdc31-2 CAN1::KANMX trp1Δ::KANMX cyh2 LYP1 ura3-1 his3-11,15 ade2-1 pURA3-CDC31*) to ~50% viability using EMS. Loss of the *pURA3-CDC31* covering plasmid was selected using 5-FOA; strains without a suppressor will spontaneously diploidize as shown in Figure 1 while those with a suppressor will remain haploid. (B) Haploid (1N) or diploid (2N) strains can be mated to a haploid to form diploid (2N) or triploid (3N) cells. The viability of meiotic products is high from diploids (88.2%, n=40 tetrads) compared to triploids (6.2%, 40 tetrads). Using this property, suppressors of diploidization were selected by mating to SLJ6750 (*MATa CDC31 can1Δ::STE2pr-HIS3MX CYH2lyp1Δ::HYGMX ura3-1 trp1-1 his3-11,15 ade2-1*) on YPD + G418 + Hyg. Following sporulation, haploid selection was carried out using SD-His-Lys-Arg+canavanine+thialysine+cycloheximide. (C) From ~100,000 EMS mutagenized cells, 61 possible suppressors were identified, and 54 were confirmed to be haploids in a secondary screen of the original mutagenized colonies by flow cytometric analysis of DNA content. Of these, 43 appeared to have mutations in the covering plasmid that allowed for growth. The remaining 11 suppressors were analyzed by tetrad dissection to ensure that suppression segregates 2:2 through at least two crosses to SLJ6121 (*MATa cdc31-2 can1Δ::STE2pr-HIS3MX TRP1 CYH2 ura3-1 his3-11,15 ade2-1 pURA3-CDC31*). An example of flow cytometry data from one hit is shown. (D) Coverage ratio of all 16 yeast chromosomes in the haploid suppressors (*ems7*, *ems9*, or *ems11*) relative to the diploid control (*EMS7*, *EMS9*, *EMS11*). Other single nucleotide polymorphisms and insertions/deletion polymorphisms identified in the haploid suppressors are listed in Table S1. (E) Quantitative PCR was performed on all 11 suppressors to determine the mean copy number of all 16 chromosomes relative to a wild-type, with chromosome XV plotted in red. Error bars, standard deviation from the mean.

It is unclear why the mutant cells are able to divide as stable diploids without forming tetraploids. One possibility is that tetraploids do indeed form but are quickly reverted to diploids through chromosome loss although the perfect ploidy of the *cdc31-2* diploids argues against this possibility. The IPL phenotype of *cdc31-2* and other mutants is thought to arise from a monopolar cell division in haploids, resulting in an aploid cell that dies and a diploid cell that is able to undergo successful divisions as a diploid (Jaspersen, Giddings, and Winey 2002; Sing et al. 2018). Additionally, acquisition of suppressor mutation that bypasses the SPB defect caused by the original mutation may be involved in diploidization (Figure 1B).

### Suppressors of cdc31-2 spontaneous diploidization

To determine the mechanisms that drive the *cdc31-2* IPL phenotype and prevent further increases in ploidy, we developed a forward genetic screen to identify suppressors of the spontaneous diploidization observed in *cdc31-2* mutants (Figure 2A). Because *cdc31-2* is a recessive mutation, we maintained cells as haploids using a plasmid containing a wild-type copy of *CDC31* (*pURA3-CDC31*), known as a covering or complementing plasmid. Haploid *MATα cdc31-2 pURA3-CDC31* cells were mutagenized with ethyl methanesulfonate (EMS), individual mutagenized cells were selected, and the covering plasmid then was removed by growth on 5-fluoroorotic acid (5-FOA). Cells that contain an IPL suppressor are haploid while the remainder spontaneously diploidize due to the *cdc31-2* allele that is uncovered following plasmid loss (Figure 2A).

In budding yeast, the ability to sexually reproduce is not controlled by chromosome number but rather by the mating type locus (MAT) present on chromosome III. Typical diploids are heterozygous for MAT (*MATa/MATα*) and are therefore able to undergo meiosis to produce four viable haploid progeny known as spores. Triploid meiosis (*MATa/MATa/MATα* or *MATa/MATα/MATα*) is catastrophic because few spores contain chromosome combinations compatible with life. In our strain background, the viability of meiotic progeny from diploid meiosis in yeast homozygous for *cdc31-2* is 88.2% while the viability of progeny from triploids is 6.2% (Figure 2B). We therefore screened for suppressors of IPL through a selection scheme involving a non-mutagenized *MATa cdc31-2 pURA3-CDC31* strain mated with the EMS-induced mutant library. In this system, only *cdc31-2* cells that contain a suppressor of spontaneous diploidization will mate to form diploids, undergo a successful meiosis and generate viable progeny. In contrast, cells that spontaneously diploidized will mate to form triploids, which will die under the meiotic selection conditions (Figure 2B).

Of ~100,000 EMS mutagenized cells that were screened as described, we isolated 61 possible suppressors of the *cdc31-2* IPL phenotype (Figure 2C). To confirm that these cells had suppressed diploidization, we performed flow cytometry on all 61 hits. Of these, 54 displayed predominately 1N and 2N peaks characteristic of haploid cells and these were pursued further. Our system utilizes selection with 5-FOA, which is converted into a toxic metabolite in yeast containing *URA3* (Boeke et al. 1987). If a mutation is introduced into the *URA3* gene by EMS, haploid *cdc31-2* cells containing the covering plasmid would be able to grow on this counter-selection. To remove these potential false positives, we tested our 54 potential suppressors and found that 43 retained the covering plasmid. Further evaluation of these remaining 11 suppressor strains showed that each was linked to a single locus in the nuclear genome. Suppression of the *cdc31-2* IPL in one tetrad is shown in Figure 2C.

From these eleven suppressors, we chose three for further characterization by Illumina sequencing. Pooled genomic DNA from 20 meiotic progeny with and without the suppressor was analyzed. No single nucleotide or insertion/deletion polymorphisms were shared among all three strains, suggesting that suppression was not caused by a change in a single gene shared by all three mutants (Table S1). An obvious suppressor mutation within the genome also was not obvious. Strikingly, all three suppressors showed increased read depth for the entire length of chromosome XV relative to control when compared to all other chromosomes (Figure 2D). Thus, all three suppressors of *cdc31-2* IPL contained two complete copies of chromosome XV while retaining a single copy of all other chromosomes.

Using a quantitative PCR assay (Wang et al. 2002), we determined that the increased copy number for chromosome XV was present in all eleven isolated suppressors (Figure 2E). Most contained a single copy of chromosomes I-XIV and chromosome XVI, however, one (*ems5*) had a more complex karyotype that may be due chromosome rearrangements such as diploidization followed by chromosome loss (Figure 2E). Alternatively, this mutant may exhibit cell to cell variation in chromosome content. Because all cells contained two copies of chromosome XV, this phenotype was further characterized as it suggests that disomy for chromosome XV can suppresses IPL of *cdc31-2*.

### Increased cdc31-2 dosage suppresses diploidization

Cells disomic for chromosome XV exhibit a number of phenotypes, including a short delay in G1 phase of the cell cycle and a small increase in cell volume compared to normal haploid cells (Torres et al. 2007; Torres et al. 2010; Bonney, Moriya, and Amon 2015). However, these phenotypes seem unlikely to be related to the mechanism of *cdc31-2* suppression since we did not recover disomies for other chromosomes with similar effects on cell size and the cell cycle, nor did they suppress *cdc31-2* IPL when directly tested. The specificity for chromosome XV suggests that suppression is linked to a gene or genes located on that chromosome, which are known to be upregulated in disomic strains relative to the rest of the haploid genome (Mulla, Zhu, and Li 2014). It seems likely that doubling the dosage of the gene(s) on chromosome XV is sufficient to alleviate the defect in SPB duplication that occurs in haploid *cdc31-2* mutants.

Importantly, the *CDC31* locus is located on the right arm of chromosome XV, making *cdc31-2* itself a leading candidate for dosage-mediated suppression.

To test the idea that *cdc31-2* itself suppresses IPL, we first tested if *cdc31-2* was necessary for suppression in the disomic strain (*cdc31-2 2xChXV*) (Figure 3A). Deletion of one copy of *cdc31-2* (*cdc31-2 2xChXV(cdc31Δ::KANMX*)) reduced growth at 23°C compared to the disomic control to a level similar to *cdc31-2* mutants (Figure 3A, C). These cells also spontaneously diploidized like *cdc31-2* (Figure 3B). Thus, it appears that the *cdc31-2* locus is necessary for suppression of IPL in the disomic strain.

**Figure 3.**
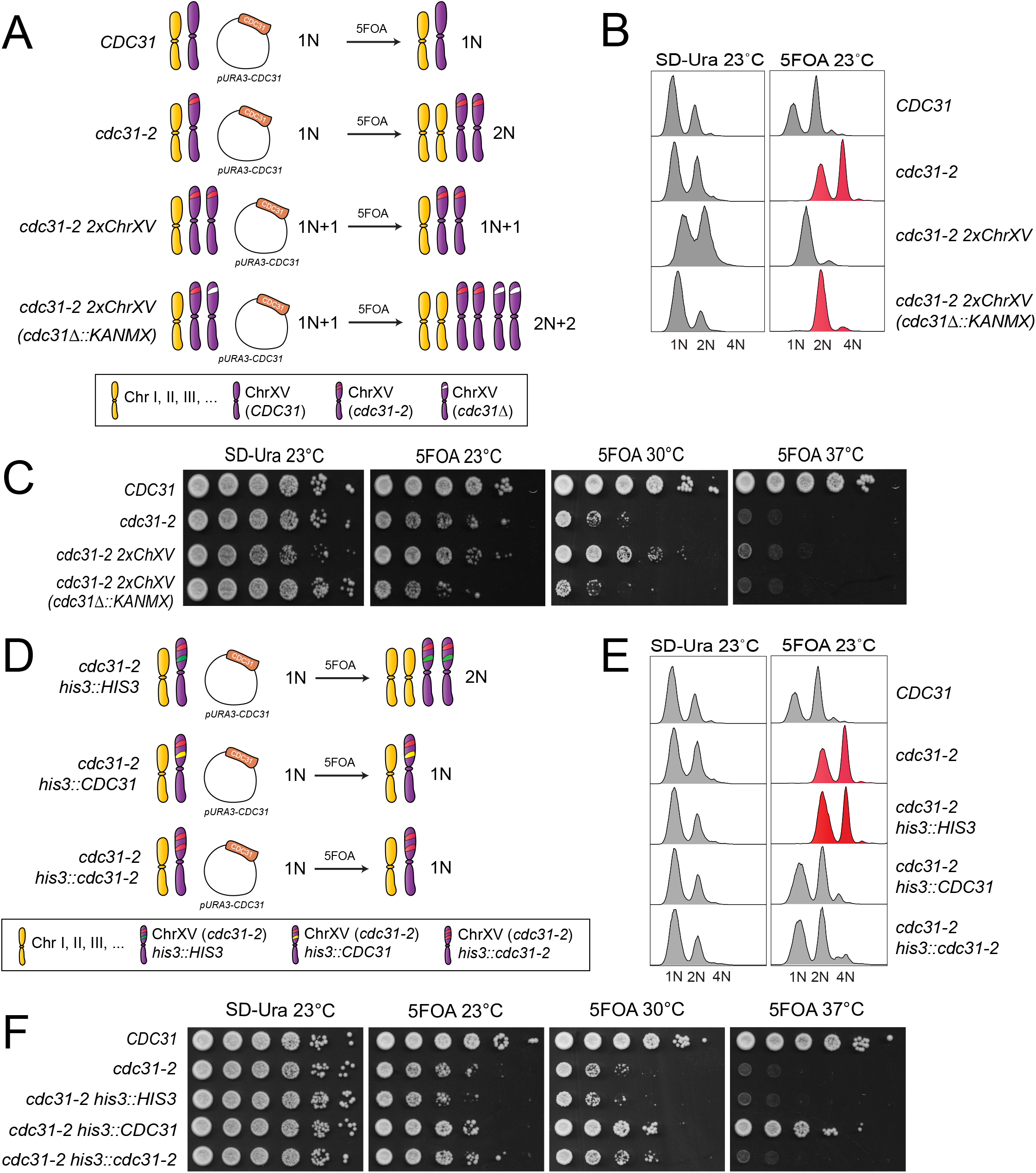
An extra copy of the *cdc31-2* gene is necessary and sufficient to suppress IPL. (A) To test if an extra copy of *cdc31-2* is necessary to suppress IPL, one copy of the *cdc31-2* locus was deleted in cells a chromosome XV disome homozygous for *cdc31-2*, was illustrated in the schematic. (B-C) The DNA content (B) and growth (C) of wild-type (SLJ7249), *cdc31-2* (SLJ809), the chromosome XV *cdc31-2* disome (SLJ7106, *cdc31-2 2xChXV(cdc31-2)*) and the deletion ((SLJ7111, *cdc31-2 ChXV(cdc31-2Δ::KANMX)*) that contain *pURA3-CDC31* were compared after growth in SD-Ura or 5-FOA at the indicated temperatures. (D) To test if an extra copy of *cdc31-2* is sufficient to suppress IPL, one additional copy of *cdc31-2* was inserted into the *HIS3* locus on chromosome XV. (E-F) The DNA content (E) and growth (F) of wild-type (SLJ7249), *cdc31-2* (SLJ809) and *cdc31-2* with an empty vector, wild-type, *CDC31* or *cdc31-2* at *HIS3* (SLJ13092, SLJ13093 or SLJ13094) were analyzed after growth in SD-Ura or 5-FOA at the indicated temperatures.

To test if an extra single copy of *cdc31-2* is sufficient to suppress IPL, we integrated a single copy of *cdc31-2* containing its endogenous promoter, terminator and coding sequence into a covered haploid *cdc31-2* strain at the *HIS3* locus on chromosome XV (Figure 3D). We also constructed isogenic strains containing an empty vector or wild-type *CDC31* at *HIS3* as controls (Figure 3D). As shown in Figure 3E, a single extra copy of *cdc31-2* integrated into the genome in combination with *cdc31-2* at the genomic locus (*cdc31-2 his3::cdc31-2-HIS3*) is sufficient to suppress IPL observed in *cdc31-2* mutants after removal of the covering plasmid on 5-FOA. These cells grow as well, or better, than *cdc31-2* mutants that spontaneously diploidize or *cdc31-2* carrying a chromosome XV disome (Figure 3C & F). This could be due the fact that a single gene rather than multiple genes is altered, resulting in little or no changes in the overall cellular proteome. Despite increased growth at 23°C and 30°C, two copies of *cdc31-2* are unable to suppress temperature sensitivity at 37°C (Figure 3C & F). This finding suggests that the copy number-based suppression of IPL at 23°C occurs via a different mechanism than dosage suppression of *cdc31^ts^* alleles at 37°C, which is probably based on bypassing, stabilizing or upregulating the mutant protein.

### SPB scaling along with ploidy change

One effect of polyploidy is an increase in the size of intracellular structures. Based on EM analysis of SPB structure, SPB diameter (measured at the SPB core) increases from 110 nm in haploid cells to 160 nm in diploids; however, SPB height does not change (Byers and Goetsch 1974) (Figure 4A-B). The length of the half-bridge also does not change when ploidy is increased, and its width may only marginally increase (Figure 4B). One possibility linking spontaneous diploidization to a subset of SPB mutants such as *cdc31-2* is based on ‘dosage suppression’ – diploids have two gene copies, and presumably twice the amount of protein, to make the same structure that is needed in both haploids and diploids. For other SPB mutants, diploidization would not have the same effect as a larger structure requiring more protein is built.

**Figure 4.**
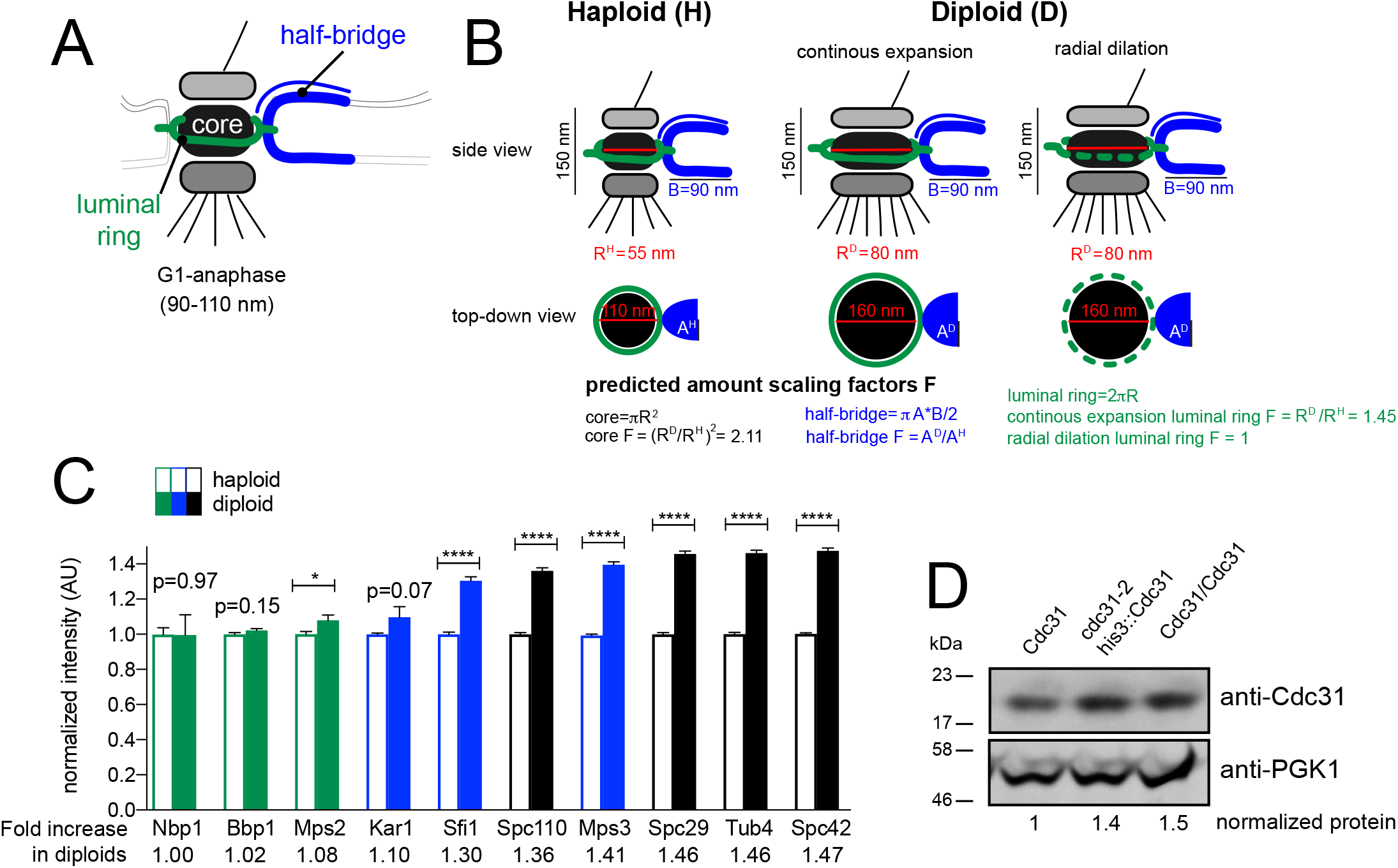
Scaling of SPB components with ploidy. (A) Schematic of the SPB showing the location of the core, luminal ring and half-bridge. (B) Side and top-down views of the SPB from haploids and diploids along with dimensions reported from EM measurements. Assuming the SPB is round and the bridge elliptical and limited to a single protein layer, theoretical scaling factors can be calculated. Based on dimensions calculated from EM measurements, which are shown, a single layer of protein in the SPB core would be 2.11 times larger in diploids. The half-bridge is thought to be a monolayer of constant length in both haploids and diploids, however, its width may scale. Two potential models for scaling of the luminal ring are depicted: a continuous scaling, where components increase proportionally to the circumference of the SPB core (1.45-fold); or radial dilation, where the amount of components do not increase. (C) Levels of fluorophore tagged protein derivatives expressed from endogenous loci in haploids or homozygous diploids were determined by quantitative imaging. For each protein, levels in haploid cells were normalized to 1. Errors, SEM with N>300 for each sample. (D) To determine levels of Cdc31, western blotting was performed using anti-Cdc31 antibody. Pgk1 served as a loading control.

A rigorous test of this idea involves the comparison of protein levels at the SPB for different components. If our hypothesis is correct, levels of core proteins will be higher in diploids compared to haploids, as the size of the core, in theory, scales approximately two-fold between haploids and diploids. Alternatively, a similar increase in ploidy should not affect levels of half-bridge proteins since its size is thought to be similar in haploids and diploids (Kilmartin 2003). We anticipated that levels of the luminal ring would also increase in size in diploids, although given the differences in geometry and organization the luminal ring may not increase to the same extent as the core (Figure 4B). To directly test these hypotheses, we compared the intensity of multiple SPB components endogenously-tagged with mTurquoise2 at the C-terminus in both haploid and homozygous diploid strains, with the exception of Kar1, which was tagged at the N-terminus. We were unable to functionally tag Cdc31, similar to previous reports (Yoder et al. 2003; Jaspersen, Giddings, and Winey 2002; Kilmartin 2003). SPB components clustered into three groups based on the intensity increase observed in diploids: no/mild (up to 1.1-fold), modest (1.1-1.4-fold) and major (over 1.4-fold) increase. Somewhat unexpectedly, none showed the anticipated increased fluorescence intensity based on scaling models and EM measurements (Figure 4C). For example, core SPB components (Spc110, Spc29, Tub4, Spc42), which underwent major scaling, showed a ~1.4-1.5-fold increase in diploids.

The group that showed a mild increase in diploids contained components of the luminal ring: Nbp1, Bbp1 and Mps2 (Figure 4C). The observation that this SPB substructure does not scale as predicted in our theoretical model could be caused by heterogeneity in ring shape (Chen et al. 2019) or by a difference in the actual mechanism of ring expansion. During post-mitotic nuclear pore complex (NPC) assembly, the membrane ring expands through a process known as radial dilation (Shahin et al. 2005; Otsuka et al. 2018). A constant total amount of protein is spread over an expanding NPC core, decreasing protein area with the increase in size. Our data suggests that the luminal ring expands by a similar radial dilation process (Figure 4C).

The levels of half-bridge proteins Sfi1 and Kar1 showed a modest increase in diploids (Figure 4C). This suggests that the width of the bridge scales a small amount from haploids to diploids. Mps3 also showed an increase; as a dual component of the half-bridge and the luminal ring (Chen et al. 2019), this is consistent with bridge scaling (as seen for Sfi1 and Kar1) and radial dilation (as seen for Nbp1, Bbp1 and Mps2). Although we were unable to visualize Cdc31 microscopically, we found that an extra copy of *CDC31* resulted in a 50% increase in total Cdc31 levels, measured by western blotting of whole cell extracts (Figure 4D). As Cdc31 is present at multiple structures throughout the cell (Fischer et al. 2004; Chen and Madura 2008; Myers and Payne 2017), this would likely reflect a moderate increase in its levels at the SPB, similar to the trend we observed for the other half-bridge components, Sfi1, Kar1 and Mps3. Overall, our data are consistent with the idea that IPL may be linked to scaling of the SPB.

### Gene dosage as a general mechanism to suppress IPL in SPB mutants

Diploidization is not unique to *cdc31-2*, but it is also observed in other SPB mutants, including some, but not, all alleles of *MPS3*, *KAR1*, *NBP1* and *MPS2* (Table 1). The fact that the phenotype is limited to half-bridge and luminal ring SPB components may be due in part to the fact that these SPB substructures do not or only modestly scale in size (Figure 4B-C). However, the observation that it is limited to certain specific alleles suggests the dosage-dependent IPL mechanism also involves the mutant protein and its role at the SPB. For example, a common defect in SPB duplication may underly the IPL phenotype (Figure 5A).

**Figure 5.**
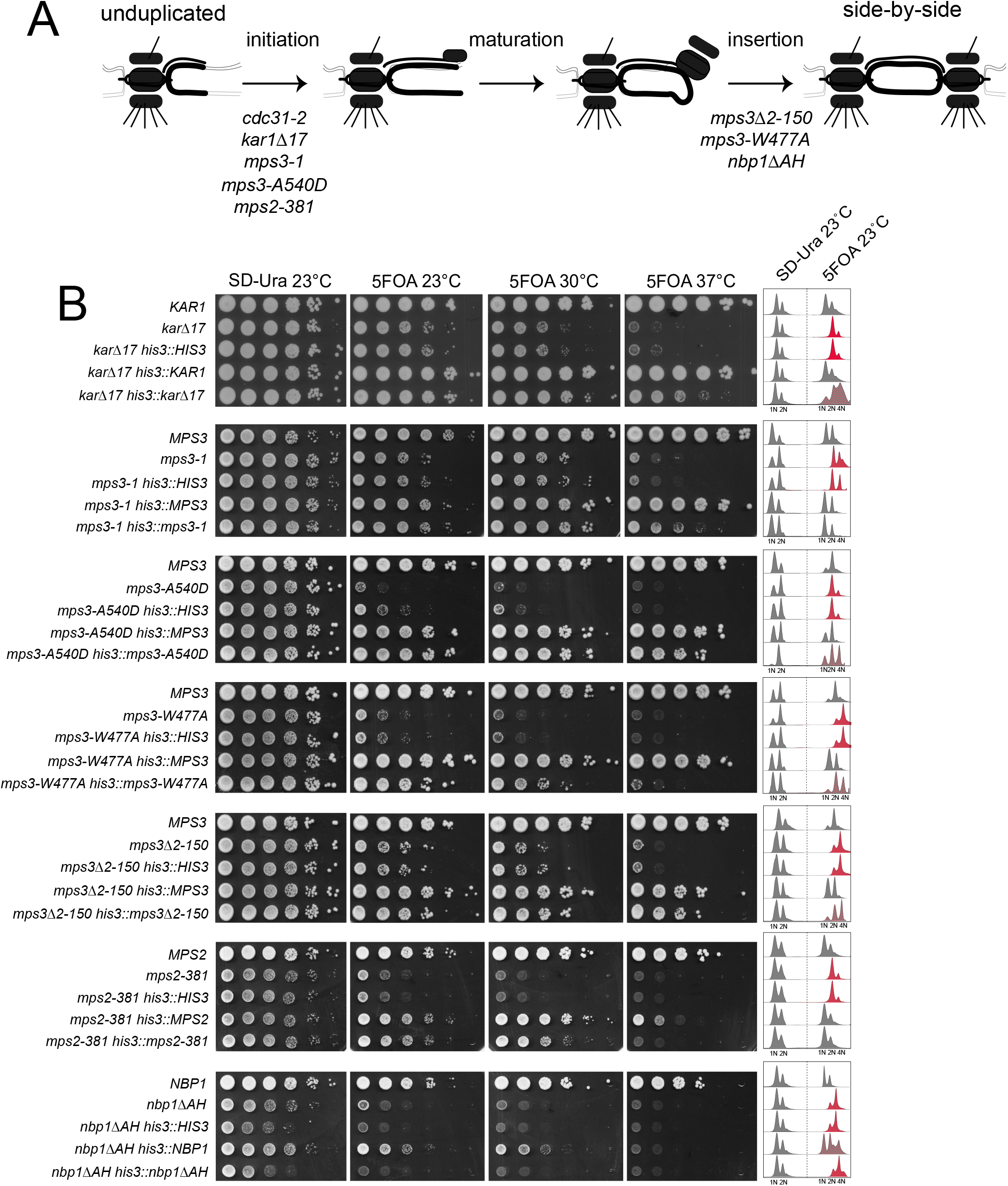
Dosage and IPL in other SPB mutants. (A) Schematic of SPB duplication pathway from an unduplicated SPB to duplicated side-by-side SPBs. Mutants defective in initiation, maturation and insertion of the new SPB have been isolated; shown are alleles required at each step that also exhibit IPL at 23°C (see Table 1). (B) To test if an extra copy of these mutant genes is sufficient to suppress IPL, one additional copy was inserted into the *HIS3* locus on chromosome XV as in Figure 3D-F. The growth and DNA content of wild-type, mutant and mutant with an empty vector, wild-type gene or mutant gene at *HIS3* were analyzed after growth in SD-Ura or 5-FOA at the indicated temperatures.

**Table 1.** Ploidy level and function of *spb ts* alleles at 23°C in W303

| SPB *ts* alleles | Ploidy at 23°C | Localization at SPB* | References |
| --- | --- | --- | --- |
| *cdc31-2* | 2N/4N | half-bridge | (Schild, Ananthaswamy, and Mortimer 1981) |
| *CDC31-16* | 1N/2N | half-bridge | (Vallen et al. 1994) |
| *kar1Δ17* | 2N/4N | half-bridge | (Vallen et al. 1994) |
| *sfi1-3* | 1N/2N | half-bridge | (Anderson et al. 2007) |
| *sfi1-7* | 1N/2N | half-bridge | (Anderson et al. 2007) |
| *mps3-1* | 2N/4N | half-bridge & luminal ring | (Jaspersen, Giddings, and Winey 2002) |
| *mps3-W477A* | 2N/4N | half-bridge & luminal ring | (Jaspersen et al. 2006) |
| *mps3-W487A* | 1N/2N | half-bridge & luminal ring | (Jaspersen et al. 2006) |
| *mps3-Y502H* | 1N/2N | half-bridge & luminal ring | (Jaspersen et al. 2006) |
| *mps3-A540D* | 2N/4N | half-bridge & luminal ring | (Jaspersen et al. 2006) |
| *mps3-F592S* | 1N/2N | half-bridge & luminal ring | (Jaspersen et al. 2006) |
| *mps3Δ2-150* | 2N/4N | half-bridge & luminal ring | (Chen et al. 2019) |
| *mps2-1* | 2N/4N | luminal ring | (Winey et al. 1991) |
| *mps2-381* | 2N/4N | luminal ring | (Jaspersen et al. 2006) |
| *ndc1-A290E* | 1N/2N | luminal ring | (Chen et al. 2014) |
| *ndc1-39* | 1N/2N | luminal ring | (Chen et al. 2014) |
| *ndc1-1* | 1N/2N | luminal ring | (Thomas and Botstein 1986) |
| *bbp1-1* | 1N/2N | luminal ring | (Schramm et al. 2000) |
| *nbp1-1* | 1N/2N | luminal ring | (Shimizu et al. 2000) |
| *nbp1-ΔAH* | 2N/4N | luminal ring | (Kupke et al. 2011) |
| *cnm67Δ* | 1N/2N | core | (Brachat et al. 1998) |
| *spc42-11* | 1N/2N | core | (Donaldson and Kilmartin 1996) |
| *spc29-3* | 1N/2N | core | (Elliott et al. 1999) |
| *cmd1-1* | 1N/2N | core | (Geiser et al. 1993) |
| *spc110-220* | 1N/2N | core | (Sundberg et al. 1996) |
| *spc97-14* | 1N/2N | core | (Knop et al. 1997) |
| *spc97-20* | 1N/2N | core | (Knop et al. 1997) |
| *spc98-2* | 1N/2N | core | (Geissler et al. 1996) |
| *tub4-1* | 1N/2N | core | (Spang et al. 1996) |

\* inner, central and outer plaque localization is denoted the core

To determine if gene dosage is a general mechanism able to suppress SPB alleles, we were interested in determining if other mutants, like *cdc31-2*, would survive as haploids if an extra copy of the mutant gene was introduced at an ectopic site in the genome. We first tested the mutant allele with the phenotype most similar to *cdc31-2*: *kar1Δ17*, which contains a partial deletion in the Cdc31 binding domain (Vallen et al. 1994). Using a covered haploid strain, we integrated a single extra copy of *kar1Δ17* into the genome of the *kar1Δ17* mutant at a marker locus. A single extra copy of *kar1Δ17* suppressed diploidization at 23°C in a small fraction of cells; however, most cells were of higher ploidy (Figure 5B). Next, we examined other SPB mutant alleles that have a SPB duplication defect similar to *cdc31-2* (Vallen et al. 1994; Jaspersen, Giddings, and Winey 2002; Jaspersen et al. 2006; Chen et al. 2019; Winey et al. 1991; Kupke et al. 2011). An extra copy of *mps3-1* or *mps2-381* completely suppressed diploidization, similar to *cdc31-2*, while *mps3-A540D* resulted in a partial rescue like *kar1Δ17* (Figure 5B). Growth fitness improved in all the alleles with doubled gene copy number at all temperatures. Although many mutants (*cdc31-2*, *mps3-1* and *mps2-381*) affecting initiation of SPB duplication can be suppressed through the addition of an extra copy of a mutant gene, increasing the dosage of other mutants only partially rescued IPL, suggesting the nature of the mutation is important to dosage-based diploidization suppression.

Examination of additional mutant alleles provided insights into the dosage-based diploidization survival mechanism. Three *MPS3* alleles (*mps3-1*, *mps3-A540D* and *mps3-W477A*) containing lesions in the C-terminal SUN domain that disrupt nucleoskeleton and cytoskeleton (LINC) complex formation exhibit different degrees of IPL suppression (Figure 5B). Interestingly, increasing the dosage of *mps3-W477A* poorly suppressed its IPL phenotype. While this could be attributed to the allele, the fact that the *mps3Δ2-150* mutant also shows a similar phenotype points to another contributing factor. Unlike most *MPS3* alleles which are defective in initiation of SPB duplication, *mps3-W477A* and *mps3Δ2-150* block SPB insertion (Figure 5A) (Friederichs et al. 2011; Chen et al. 2019). Examination of two additional insertion mutants (*mps2-1* or *nbp1ΔAH*) showed that extra copies of these genes also did not rescue IPL or restore growth when added in extra copy (Figure 5A; data not shown). Collectively, these data support the idea that spontaneous diploidization serves as a dosage suppressor for specific SPB mutants that primarily affect initiation of SPB duplication.

## Discussion

While lab strains of budding yeast are often maintained as stable haploid or diploid populations, polyploidy is common in natural yeast isolates (Ezov et al. 2006). The higher DNA state of triploid or diploid yeast allows for adaptation through the accumulation of mutations, some of which can be beneficial for fitness. However, higher ploidy is also associated with CIN, raising the question as to how cells maintain a stable ploidy.

Over the years, mutations in a number of pathways have been linked to spontaneous diploidization of haploid yeast strains, including: 1) mating type mutations, which restores the diploid state through mating-type switching (Lee et al. 2010); 2) cytokinetic mutants that fail in cell division to give rise to binucleate diploids (Rancati et al. 2008); and 3) spindle mutants that result in errors in chromosome segregation to produce mononuclear diploids (Sing et al. 2018; Luca and Winey 1998; Antoniacci et al. 2004; Chambers et al. 2012; Pinto and Winston 2000; Yu et al. 2011; Chan and Botstein 1993). Here, we investigated the IPL phenotype associated with certain mutations in the yeast SPB. Given SPB mutations are linked to chromosome segregation defects, an unresolved issue is why these alleles can be propagated as diploids under permissive conditions. While it is possible that tetraploids and octaploids do form, these higher ploidies are not detected for most alleles using flow cytometry. WGS of multiple SPB mutants also does not indicate disomes or other aneuploids are formed at least at the population level. For *cdc31-2* mutants, it was necessary to treat cells with a chemical mutagen and passage them through meiosis to create a disomic strain. This indicates that diploidization is the preferred ploidy of certain SPB mutants. Our goal was to understand the mechanism behind this ploidy regulation.

We describe how specific SPB mutations undergo dosage-based spontaneous diploidization. This phenotype is due, at least in part, to scaling of SPB architecture from haploid to diploid cells. Partial loss of function mutants that affect the first step of SPB duplication and whose gene products localize to SPB substructures that do no scale (luminal ring) or moderately scale (half-bridge) use diploidization to increase their copy number and to eliminate defects in SPB function at the permissive temperature. Thus, *cdc31-2*, *mps3-1* and *mps2-381* mutants are able to propagate as diploids. Importantly, extra copies of these genes do not overcome the growth defects and SPB phenotypes seen at higher temperatures, presumably because the mutant proteins are unstable at 37°C or the SPB duplication defect is not bypassed. Other SPB alleles such as *kar1Δ17*, *mps3-A540D*, *mps3-W477A* and *nbp1-AH* also show diploidization, but the IPL phenotype of haploids was not fully rescued by an additional copy of the mutant gene. While our ectopic rescue constructs may not express the gene at levels comparable to the endogenous locus, the fact that a disome did not suppress the IPL associated with *mps3-A540D* suggests a non-dosage dependent diploidization rescue pathway also exists.

In previous work, incompatible allometry within the spindle was linked with the CIN phenotype in *S. cerevisae* and other fungi (Jelenic et al. 2018; Hoyt, Stearns, and Botstein 1990). Although the surface area of the SPB increases to expand microtubule nucleation capacity, the length of the pre-anaphase spindle does not change in tetraploids compared to diploids even though tetraploid cells have twice the DNA content. As a result, the incidence of syntelic (monopolar) chromosome attachments is higher in tetraploids (Storchova et al. 2006). The loss of chromosomes from tetraploid *C. albicans* is so dramatic that it results in diploid or near diploid progeny (Bennett and Johnson 2003). Unlike the SPB core, we show here that the luminal ring does not increase in size to a similar extent in diploids compared to haploids. This finding strongly suggests that, similar to the NPC, this region of the SPB expands and contracts via radial dilation. This mechanism of scaling could facilitate cell cycle changes in the luminal ring size without the need to incorporate more protein. For example, in haploids the SPB core size increases from 90 nm in G1 to 110 nm in mitosis (Byers and Goetsch 1974); radial expansion of the luminal ring would accommodate this increase without incorporation of new protein. Similarly, as SPB size decreases back to 90 nm upon anaphase exit, radial contraction of the SPB would allow the luminal ring to shrink by increasing protein density.

In metazoans, centrosome function is regulated by factors involved in its duplication, maturation and microtubule nucleation. Centrosomal defects are linked to errors in chromosome segregation, in part due to the role of centrosomes in spindle organization. However, centrosomes also contribute to the size of the mitotic spindle. Our work further illustrates another possible role for centrosomes in driving genetic changes – the acquisition of mutations in genes such as centrin, which is conserved throughout eukaryotes, might promote stable genome amplification, including the genome duplications seen throughout evolution in fungi and metazoans or during tumorigenesis in humans.

## Materials & Methods

### Yeast strains and plasmids

All strains are derivatives of W303 (*ADE2 trp1-1 leu2-3,112 ura3-1 his3-11,15 can1-100 RAD5+*) and are listed in Table S2. Standard techniques were used for DNA and yeast manipulations, including C-terminal tagging with fluorescent proteins and gene deletion by PCR-based methods (Gardner and Jaspersen 2014). Single copy integrating plasmids containing SPB genes were made by PCR amplifying the open-reading frame, ~700 bp of promoter sequence and ~200 bp of the terminator from genomic DNA and assembling this DNA into pRG203MX using Gateway assembly (Gnugge, Liphardt, and Rudolf 2016). Mutations were introduced by site-directed mutagenesis of the wild-type gene using the QuikChange mutagenesis kit (Agilent). Sequencing was performed to confirm correct base pair substitutions or deletions were made.

### Screen for cdc31-2 IPL suppressors

SLJ6749 (*MATα cdc31-2 CAN1::KANMX trp1Δ::NATMX cyh2 LYP1 ura3-1 his3-11,15 ade2-1 pURA3-CDC31*) was grown overnight at 30°C in SC-Ura plus casamino acids to an OD_600_ of ~2.0. Cells were harvested and individual aliquots were mutagenized with a dosage of EMS that resulted in ~50% lethality compared to an untreated control. Following mutagenesis, cells were plated to YPD at 23°C at which time individual colonies were cherry-picked into 96-well plates to allow for automated pinning using the Singer ROTOR (Singer Instruments). Next, loss of the *pURA3-CDC31* covering plasmid was selected by growing cells on 5-FOA for 3 d at 23°C. Mating to SLJ6750 (*MATa cdc31-2 can1Δ::STE2pr-HIS3MX CYH2 lyp1Δ::HYGMX ura3-1 trp1-1 his3-11,15 ade2-1*) was performed overnight on YPD; mated cells were selected by growth on YPD containing 200 μg/ml G418 and 300 μg/ml hygromycin for 3 d. Cells were transferred onto sporulation media for 3 weeks at 23°C. Meiotic progeny were selected by two rounds of growth on SD-His-Lys-Arg containing 50 μg/ml canavanine, 50 μg/ml thialysine and 10 μg/ml cycloheximide for 3 d at 23°C. Suppressors of *cdc31-2* IPL give rise to colonies under these growth conditions.

From ~100,000 EMS mutagenized cells, 61 possible suppressors were identified. Flow cytometric analysis of DNA content showed that 54 exhibited peaks at 1N and 2N, which are typically observed in haploid yeast. Of these, we found using plasmid rescue, PCR and sequencing that 43 contained mutations in the *URA3* gene on the covering plasmid that allowed for growth on 5-FOA, thus these were false positives. The remaining 11 suppressors were analyzed by tetrad dissection to ensure that suppression segregates 2:2 through at least two crosses to SLJ6121 (*MATa cdc31-2 can1Δ::STE2pr-HIS3MX TRP1 CYH2 ura3-1 his3-11,15 ade2-1 pURA3-CDC31*).

### Whole genome sequencing (WGS)

Using 20 four-spored tetrads from a cross between an EMS-induced hit and SLJ6121, we identified the two progeny from each tetrad that were diploid (control) and the two progeny from each tetrad that were haploid (and therefore contained an ems hit). To ensure equal representation of colonies, each was individually grown, normalized by OD_600_, then mixed to achieve equal number of all 40 cells in the control and ems hit pools. Genomic libraries were prepared using the Illumina Mate Pair library kit and prepared for paired-end sequencing on the Illumina MiSeq as previously described (Birkeland et al. 2010). Downstream sequence analysis was performed (Birkeland et al. 2010). Reads were aligned to sacCer3 using bwa version 0.7.15-r1140 (Li and Durbin 2009) and single nucleotide polymorphisms (SNPs) and insertion/deletion polymorphisms were identified using SAMtools version 1.5 (Li et al. 2009). SNP and insertion/deletion polymorphisms were annotated using snpEff version 4.3 (Cingolani et al. 2012). Coverage was calculated using BEDTools version 2.25.0 (Quinlan and Hall 2010). Results are listed in Table S1.

### Flow cytometry and qPCR karyotyping

DNA content was analyzed by flow cytometry in sonicated cells that had been fixed with 70% ethanol for 1 h at room temperature, treated with RNAse (Roche, Basel, Switzerland) and Proteinase K (Roche) for 2 h to overnight at 37°C and stained with propidium iodide (Sigma-Aldrich, St. Louis) in the dark at 4°C overnight. Samples were analyzed on a MACSQuant FACS Analyzer (Miltenyi Biotec) and data was displayed using FlowJo software (Tree Star, Ashland, OR). qPCR karyotyping was performed using centromere proximal primers for each chromosome arm as previously described (Pavelka, Rancati, and Li 2010).

### Growth assay

To analyze growth phenotypes, 5 OD_600_ of cells from each strain were serially diluted in 10-fold, and ~7 μl of each dilution was spotted on SD-Ura or SD plates containing 5-FOA (Sigma Aldrich). Plates were incubated at indicated temperatures for 2-4 days.

### Image analysis

Live cell imaging was used to study spindle structure in cells containing GFP-Tub1 (microtubules) and Spc42-Cherry (SPBs) using a Perkin Elmer (Waltham, MA, USA) Ultraview spinning disk confocal microscope equipped with a Hamamatsu (Hamamatsu, Japan) EMCCD (C9100-13) optimized for speed, sensitivity and resolution. The microscope base was a Carl Zeiss (Jena, Germany) Axio-observer equipped with an αPlan-Apochromat 100x 1.46NA oil immersion objective and a multiband dichroic reflecting 488 and 561 nm laser lines. GFP images were acquired with 488 nm excitation and 500-550 nm emission. mCherry images were acquired with 561 nm excitation and 580-650 nm emission. Data were acquired using the PerkinElmer Volocity software with a z spacing of 0.4 μm. Exposure time, laser power and camera gain were maintained at a constant level chosen to provide high signal-to-noise but avoid signal saturation for all samples. Images were processed using Image J (NIH, Bethesda, MD). A representative z slice image is shown. Cells were considered to be large-budded if the bud size was >30% the size of the mother cell.

Images for SPB intensity quantification in isogenic haploids and diploids were captured with a Nikon Spinning Disk controlled with NIS-Elements Viewer software equipped EMCCD camera and a PlanApo 100x 1.4 NA objective. Parameters, including laser power, exposure time, z-spacing and number of stacks, were set to identical value. Quantitation of levels of SPB proteins was performed with custom plugins (freely available at http://research.stowers.org/imagejplugins) written for ImageJ (NIH, Bethesda, MD). Prior to analysis, raw images were processed with background subtraction and summed to form a single plan image. Individual SPBs were identified using an ImageJ internal function “Find Maxima” and then chose “single points” as output. A circular ROI with a size of 7 pixel was drawn on each single point to cover individual SPB. Integrated intensity was then calculated on all ROIs.

### Western blotting and quantification

Pelleted cells were washed in PBS and frozen in liquid nitrogen. Thawed pellets were resuspended in 1 ml lysis buffer (50 mM Tris, pH 7.5, 150 mM NaCl, 0.1%NP-40, 1 mM DTT, 10% glycerol and 1 mg/ml each pepstatin A, aprotinin and leupeptin) and ~100 μl of glass beads were added prior to bead beating for 1 min x 5 with 2 min on ice between beatings. Samples were spun at 5000 rpm for 2 min and the supernatant was transferred to a new tube. Protein concentration was determined using a NanoDrop Spectrophotometer (Thermo), and equivalent amounts of lysate were analyzed by SDS-PAGE followed by western blotting. Whole cell extracts prepared by bead beating into SDS sample buffer to determine expression levels of baits. The following primary antibody dilutions were used: 1:1000 anti-Cdc31 (in the lab) and 1:5000 anti-Pgk1 (Invitrogen). Alkaline phosphatase-conjugated secondary antibodies were used at 1:10000 (Promega). Western blot band intensity was analyzed with ImageJ Gel quantification tool.

## Acknowledgements

We thank Giulia Rancati for strains and are grateful to Scott Hawley, Sarah Smith and members of the Jaspersen Lab for helpful suggestions throughout the project and for their comments on the manuscript. Original data underlying this manuscript can be downloaded from the Stowers Original Data Repository at http://www.stowers.org/research/publications/LIBPB-1533. Fastq files associated with this project can be found at the National Center for Biotechnology Information under BioProject ID PRJNA632749. Research reported in this publication was supported by the Stowers Institute for Medical Research and the National Institute of General Medical Sciences of the National Institutes of Health under award number R01GM121443 (to SLJ). The authors declare no competing financial interests.

## Author contributions

SLJ conceived the experiments, JC, ZX and AMC constructed strains, ZX performed the screen, DMM analyzed genomic data and JC did the follow-up analysis. SM and WDB did qPCR and automated flow cytometry analysis, and ZY acquired images and wrote software for image quantitation. JC and SLJ prepared figures and wrote the paper with input from all the authors.

## Abbreviations

MTOC: microtubule-organizing center;
SPB: spindle pole body;
IPL: increase-in-ploidy;
CIN: chromosome instability;
EMS: ethyl methanesulfonate;
WGS: whole genome sequencing;
EM: electron microscopy;
NE: nuclear envelope;
NPC: nuclear pore complex;
FACS: Fluorescence-activated cell sorting;
SEM: standard error of the mean;
5-FOA: 5-fluoro-orotic acid; SNP, single-nucleotide polymorphism

## Supplementary Material

**Table S1.** Single nucleotide and insertion/deletion polymorphisms annotated with SnpEff for three *cdc31* suppressors

| Chromosome | Position | Reference | Change | VCF Score | Location | Gene | Protein Change | Type |
| --- | --- | --- | --- | --- | --- | --- | --- | --- |
| *cdc31-2 ems7* |  |  |  |  |  |  |  |  |
| XVI | 769362 | T | C | 222 | intergenic | na | na | na |
| *cdc31-2 ems9* |  |  |  |  |  |  |  |  |
| III | 307421 | G | A | 225 | intergenic | na | na | na |
| III | 307422 | C | A | 225 | intergenic | na | na | na |
| IX | 438797 | C | T | 222 | intergenic | na | na | na |
| V | 6586 | A | G | 222 | intergenic | na | na | na |
| V | 6594 | T | C | 221 | intergenic | na | na | na |
| V | 6600 | A | G | 221 | intergenic | na | na | na |
| VI | 92485 | C | T | 222 | genic | *BUD27* | V500V | synonymous |
| X | 354706 | T | C | 219 | intergenic | na | na | na |
| XIII | 587695 | T | C | 205 | intergenic | na | na | na |
| *cdc31-2 ems11* |  |  |  |  |  |  |  |  |
| XVI | 769362 | T | C | 222 | intergenic | na | na | na |
| XVI | 769434 | T | G | 222 | intergenic | na | na | na |

\* shown are variants detected in the genome of the mutant compared to control.

**Table S2.**
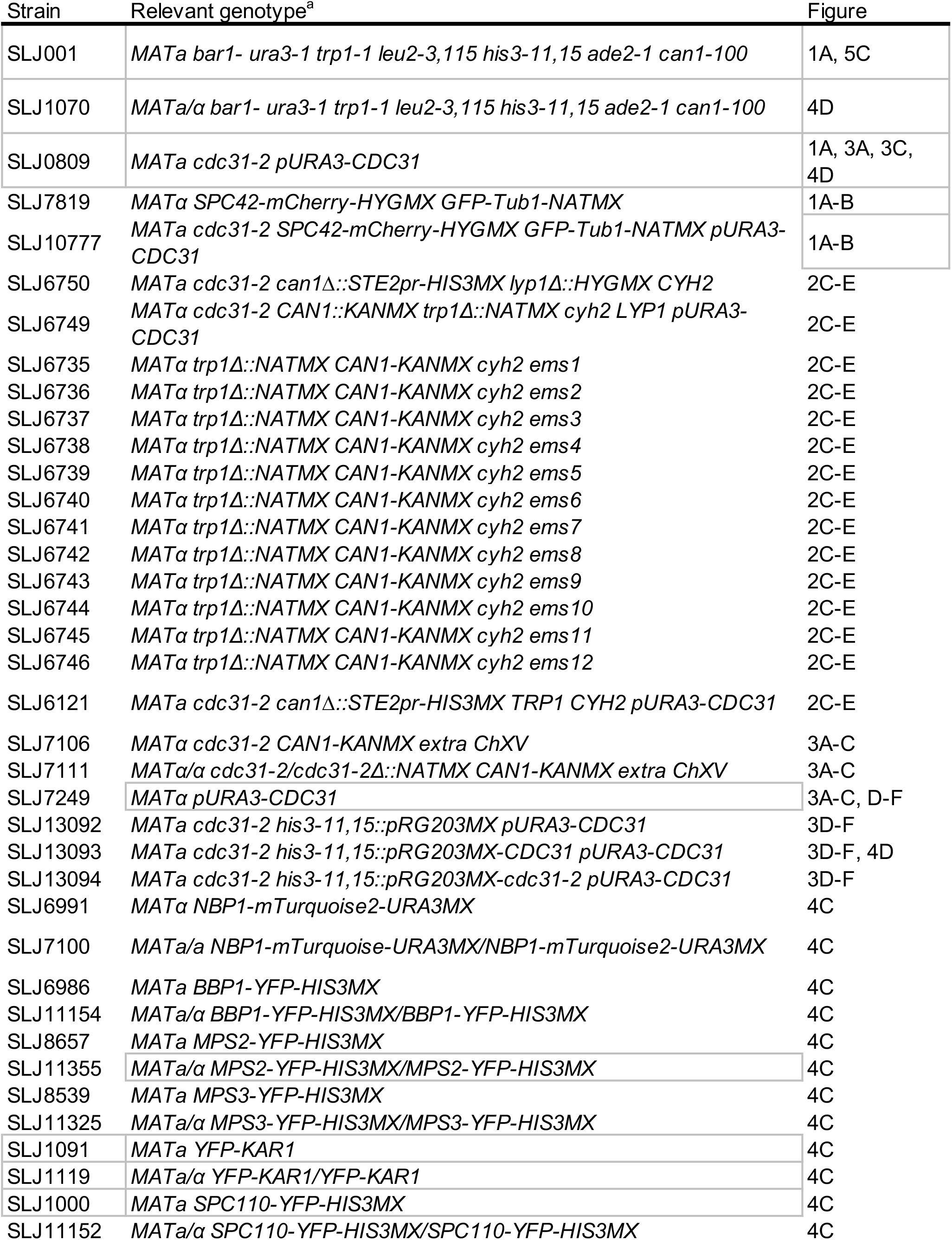
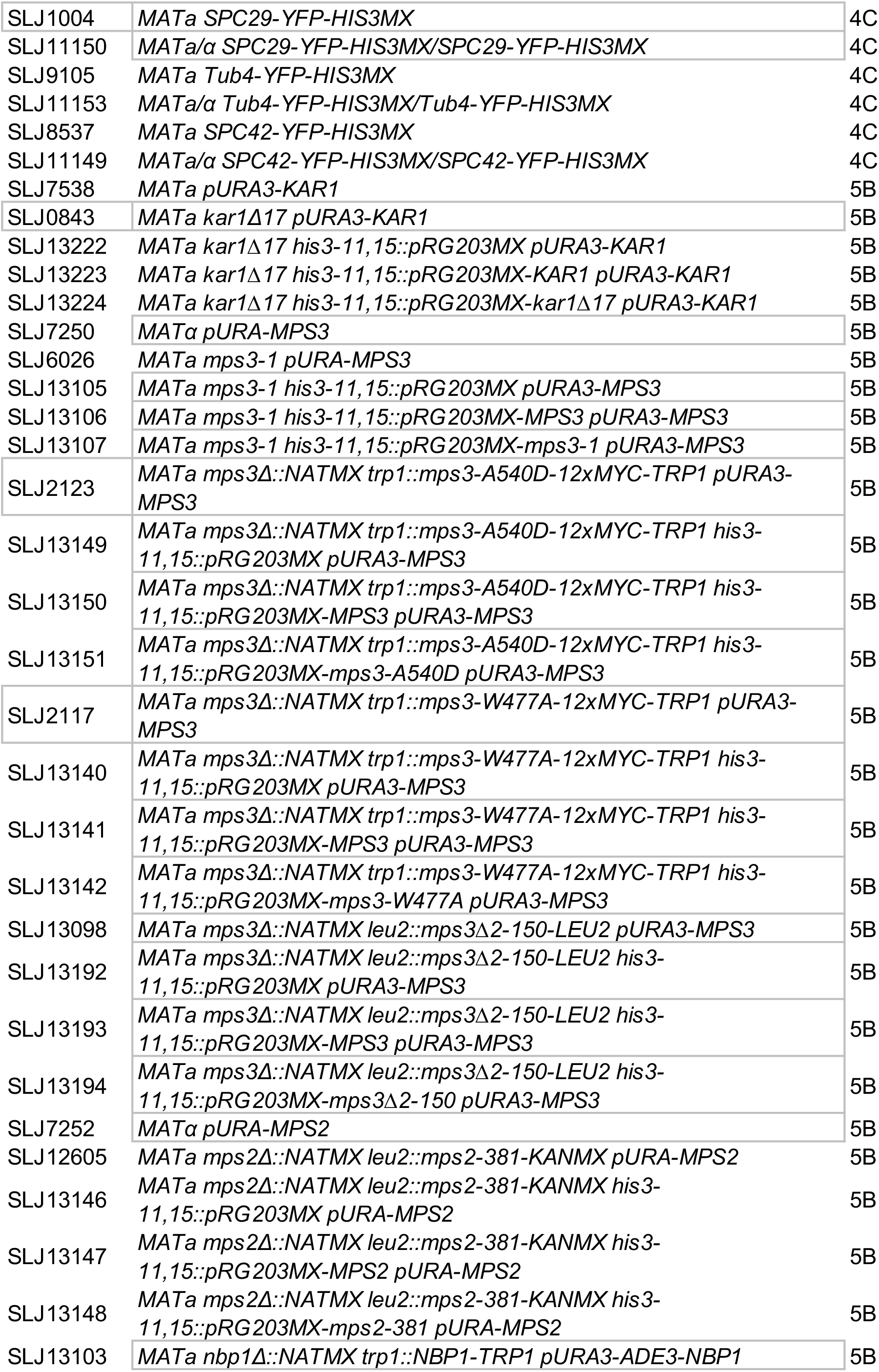
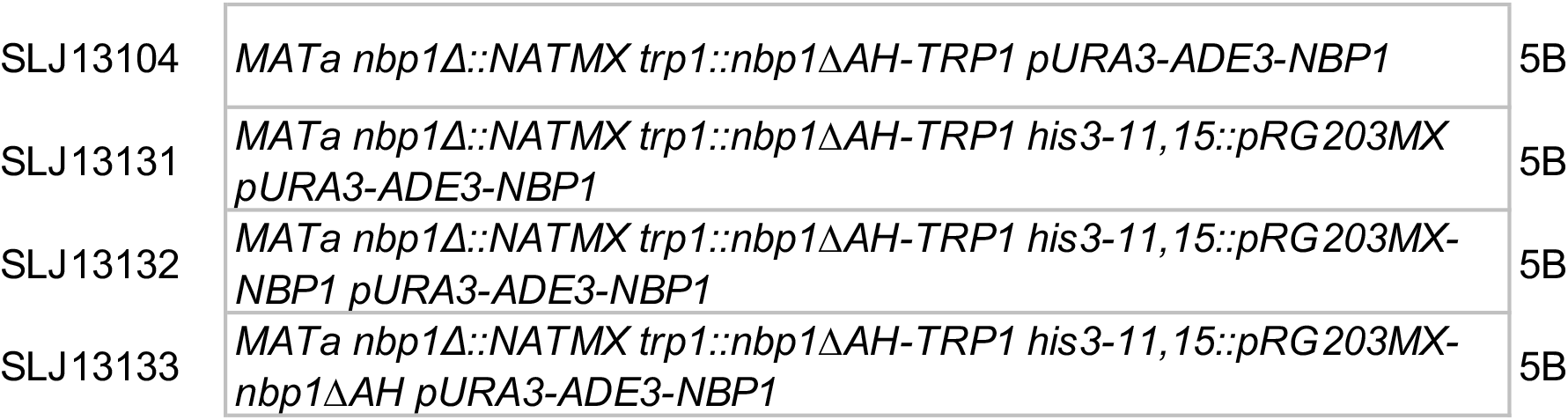
Yeast strains

